# Distinct Molecular Processes in Placentae Involved in Two Major Subtypes of Preeclampsia

**DOI:** 10.1101/787796

**Authors:** Zhonglu Ren, Yunfei Gao, Yue Gao, Guanmei Liang, Qian Chen, Sijia Jiang, Xiaoxue Yang, Cuixia Fan, Haizhen Wang, Jing Wang, Yi-Wu Shi, Chaoqun Xiao, Mei Zhong, Yanhong Yu, Xinping Yang

## Abstract

**Purpose:** Patients with preeclampsia display a spectrum of onset time and severity of clinical presentation, yet the underlying molecular bases for the early-onset and late-onset clinical subtypes are not known.

**Methods:** Since the root cause of PE is thought to be located in the placentae, RNA-seq has been performed on 65 high-quality placenta samples, including 33 from 30 patients and 32 from 30 control subjects, to search for molecular features.

**Results:** Two functionally distinct sets of dysregulated genes have been identified in two major subtypes: metabolism-related genes, notably transporter genes, in early-onset severe preeclampsia (EOSPE) and immune-related genes in late-onset severe preeclampsia (LOSPE), while the late-onset mild preeclampsia (LOMPE) is not to be distinguished from normal controls. A small number of dysregulated transcription factors are found to drive the widespread gene dysregulation in both EOSPE and LOSPE.

**Conclusion:** These results suggest that EOSPE and LOSPE have different molecular mechanisms, whereas the LOMPE may have no placenta-specific causal factors.

**DISCLOSURE:** All authors have approved the manuscripts and agreed with their submission to *Genetics in Medicine*. No part of the manuscripts has been submitted elsewhere, nor is any part under consideration for publication at another journal. There are no conflicts of interest to disclose.

## INTRODUCTION

Preeclampsia (PE) is hypertension presenting after 20 weeks of gestation combined with signs of damage to other maternal organs, mostly liver and kidney. The diagnostic criteria include systolic blood pressure higher than 140 mm Hg, and/or diastolic blood pressure above 90 mm Hg, in addition to proteinuria of over 300 mg/day.^1^ It can be classified as early- or late-onset with a gestational age cutoff of 34 weeks.^2, 3^ The global incidence rate is 4.6% of pregnancies,^4^ leading to 60,000 maternal deaths per year.^5^ The grave consequences of PE on maternal health are mainly attributed to cerebral edema, intracranial hemorrhage and eclampsia,^6, 7^ which some studies have suggested to be caused by abnormal functions of the autonomic nervous system mediated by placenta-derived factors.^8–10^ PE has significant consequences on fetal development and growth, often leading to perinatal and infant morbidity or mortality^1^ and contributes preterm births^11^ and ∼15% cases of fetal growth restriction (FGR).^12, 13^ In the long term, PE affects brain development and functions of the offspring^14^, leading to intellectual disability,^15^ epilepsy,^16^ autism^17–19^ and schizophrenia.^20, 21^

Since the symptoms of a pregnant woman with PE usually resolve after delivery and histopathological placental changes can be detected at that time, the placenta has been perceived as the root cause of PE. A widely accepted disease model for the placental origin includes two stages of PE development.^22, 23^ During the first stage, abnormal placentation in early gestation plays a role, such as poor trophoblast invasion and incomplete vascular formation of spiral arteries, which lead to placenta dysfunctions; and in the second stage, the dysfunctional placenta releases factors into the maternal blood, which in turn lead to hypertension and organ damage.^22, 23^ This model promoted intensive studies on placentae as root cause of PE, and increasing knowledge leads to finer classification of stages.^24^ However, the PE-associated molecular processes originating in the placenta remain largely unknown, and there are insufficient molecular criteria for distinction of the clinical subtypes.

Some PE cases occur in families, suggesting genetic factors play a role in its pathogenesis. First-degree relatives of women with PE have a 5-fold higher risk of developing the disease.^25^ Men born from a PE pregnancy are also at risk of fathering PE pregnancies,^26, 27^ demonstrating the placental genotype contributes to PE. The heritability of PE is about 50%.^28, 29^ Genome-wide linkage studies on pedigrees have identified risk loci on chromosomes 2p13, 2q23, 11q23, 10q22, 22q12, 2p25, 9p13, 4q32 and 9p11.^30–34^ Genome-wide association studies (GWAS) have found some variants associated with PE.^35, 36^

Patients with early-onset PE have more severe maternal and perinatal complications than those with late-onset PE.^2^ We do not know, however, if early-onset PE and late-onset PE have a common or distinct underlying pathogenesis at the molecular level. Numerous placental microarray studies failed to identify distinct molecular processes, due to small sample sizes or insufficient grouping of clinical subtypes.^37–44^ Some analyses use no classification of samples^38, 43^ and some focus on severe PE,^44^ or early-onset severe PE.^45^ Although several microarray studies contain both early- and late-onset PE placental samples, these studies investigated differential expression between all PE samples and controls, or between early and late PE,^39, 41, 42^ but did not investigate differential expression in subtypes by comparing them to normal controls. A recent RNA-seq study, done on placentae of 9 PE samples (in three pools) and 9 controls (in three pools), found 53 PE-associated genes.^46^ Although these studies point to placental mechanisms underlying PE involved in angiogenesis,^39, 42^ immune functions^37^ and hypoxia,^44^ it is still not clear whether the molecular pathways involved in the functional defects of placentae are different in clinical subtypes of this complex disorder. Moreover, if transcriptomes are compared between more homogeneous samples of each particular subtype and normal subjects, more complete sets of dysregulated genes might be revealed for each subtype.

In order to systematically search for placental molecular signatures associated with clinical subtypes, we carried out transcriptome sequencing on 65 high quality placenta samples, including 33 from 30 PE patients and 32 from 30 normal subjects, and found dysregulated genes associated with two distinct sets of pathways involved in two the major clinical subtypes.

## MATERIALS AND METHODS

### Patients

We collected the tissue samples at the Department of Obstetrics & Gynecology of Nanfang Hospital in China from January 2015 to July 2016. The clinical characteristics of each patient were extracted from the medical records, which strictly followed the American Board of Obstetrics and Gynecology, Williams Obstetrics 24th edition. Samples of placental tissues were collected from the mid-section placental tissues of the placenta, between the chorionic and maternal basal surfaces, at four different positions within 5 minutes after delivery. These placental tissues were placed into RNAlater® solution and stored at −80 °C. This research has been approved by the Ethnics Board of Nanfang Hospital of Southern Medical University, and all patients have signed the informed consent.

Preeclampsia (PE) is characterized by new-onset hypertension (≥140/90 mmHg) and proteinuria at ≥20 weeks of gestation, or in the absence of proteinuria, hypertension with evidence of systemic disease.^47^ If patients with PE have systolic blood pressure ≥160 mmHg or diastolic blood pressure ≥110 mmHg, or other organ failures, we call them “preeclampsia with severe features” or briefly “severe PE”. According to gestational age at diagnosis or delivery, PE can be divided as early-onset (< 34 weeks) and late-onset (≥ 34 weeks).^48, 49^ Although other gestational age cut-offs have been suggested, 34 weeks remains the most commonly used,^50, 51^ presumably as the rate of neonatal morbidities declines considerably after reaching this time point.

According to above criteria and clinical record of each case, we grouped our PE patients into three clinical subtypes: late-onset mild PE (LOMPE), early-onset severe PE (EOSPE) and late-onset severe PE (LOPSE). The clinical information of the three clinical subtypes and comparisons between the subtypes were listed in Table S1.

### RNA-seq

Total RNA was isolated using the RNeasy® Plus Universal Mini Kit (Qiagen) according to the manual protocol. RNA-seq was carried out at Berry Genomics Corporation (Beijing, China). Briefly, RNAs with polyA tails were isolated, and double-stranded cDNA libraries were prepared using TruSeq RNA Kit (Illumina) followed by paired-end sequencing using Illumina Hiseq 2500. In order to computationally remove the cord blood contamination, RNA-seq was also done on two samples of cord blood. The reads from the clean data of RNA-seq were aligned to the human reference genome (hg38) with STAR^52^ and counted using HTSeq^53^ with union mode. Raw counts for annotated genes (protein coding genes, rRNA, microRNA, LncRNA, pseudogenes and so on) in the General Transfer Format (GTF) annotation file were obtained. The reference genome sequence and GTF annotation files were downloaded from the Ensembl website (http://asia.ensembl.org/index.html) in the alignment and counting reads steps.

Annotation information, such as gene location, gene types, gene symbols, for differential expression genes were extracted from EnsDb.Hsapiens.v86 package. Four brief categories of gene biotypes were divided following the table listed on the GENCODE website page (https://www.gencodegenes.org/gencode_biotypes.html). Gene Ontology terms overrepresentation, Disease Ontology, Pathway and functional annotation analyses were used clusterProfiler^54^ and DOSE^55^ packages. All analyses were done in R-platform, and all packages and public datasets were listed in **Table S9**.

### Computational removal of blood contamination

The contaminated cord blood cells could not be totally washed from the placenta tissue samples, so the extracted RNAs contained some from cord blood cells. To better study the difference in gene expression of placenta tissues between PE patients and controls, we removed the contamination using the expression level of a cord blood marker gene for normalization. Cord blood samples from two normal pregnancy women were sequenced using the same RNA-seq method. Differentially expressed gene analysis was applied to two cord blood samples against two normal placenta samples to identify marker genes for cord blood cells. We used the Hemoglobin Subunit Mu (HBM), a subunit of hemoglobin that is specifically expressed in red blood cells, and recalibrate the raw count of genes accordingly in the following equation:

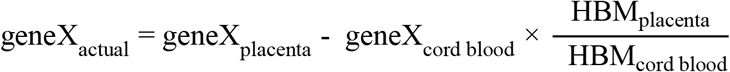

The raw counts of genes in each sample before and after removing blood contamination were showed in **Table S2**.

### Clustering analysis of PE samples only

The EOSPE samples were separated from other samples in clustering, whereas LOMPE and LOSPE samples were clustered together with normal samples (Fig. 1B). To explore clusters of PE samples, we performed clustering analysis on PE samples only, including EOSPE, LOSPE and LOMPE samples, excluding normal samples. The top 500 most variances (genes) across all PE samples were used for clustering (**Figure S1B**). All of the LOMPE samples and most of the LOSPE samples clustered together on the left side of the graph as cluster-1, whereas all EOSPE samples clustered on the right side as cluster-2, together with the four LOSPE samples.

**Fig. 1.**
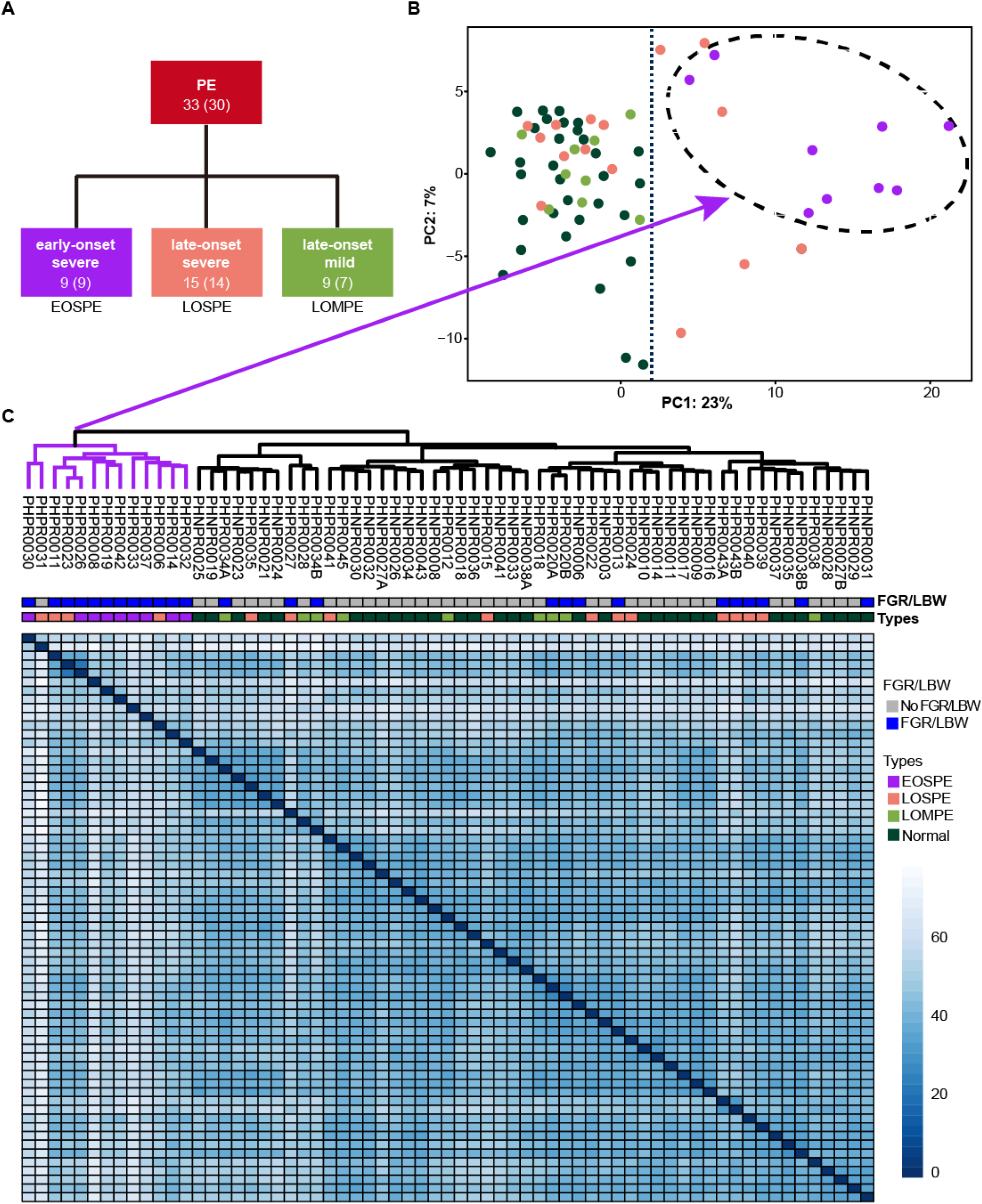
Clinical subtypes and sample clustering using RNA-seq data. **(a)** Clinical subtypes of PE patients: early-onset severe PE (EOSPE), late-onset severe PE (LOSPE), late-onset mild PE (LOMPE). The numbers under each subtype are the numbers of placentae (not in brackets) and the numbers of patients (in brackets). (**b)** Principal component analysis (PCA). The top two principal components in this dataset represent up to 30% of all variations. Samples of EOSPE clustered on the right side (within the dashed circle), while samples of LOSPE and LOMPE are clustered together with normal samples on the left side. (**c)** Heatmap of sample-sample distance clustering using the Ward’s method. The EOSPE samples are grouped together as the branch on the left, roughly corresponding to the samples in the right area of the PCA plot. The LOSPE and LOMPE samples are clustered together with normal samples on the right side. Only four LOSPE samples were clustered with EOSPE.

In addition, we carried out a comparison of the clinical characteristics of the patients between the two clusters and detected significant differences in blood pressure, gestational age of delivery, baby weight, FGR or LBW and proteinuria (**Table S1**). Of the 14 babies (13 placenta samples) in subclass-2, 12 babies (85.7%) had FGR or LBW, whereas 10 of 20 babies (50%) had FGR or LBW in subclass-1. The mean length of gestation in two clusters were 264 (37.7 weeks) and 228.8 (32.5 weeks), and patients in the two clusters showed different weeks at onset of PE. The quantification of proteinuria in the two clusters also had significant differences. In other words, PE samples in cluster-1 showed late-onset, relatively moderate symptom, and less harmful outcomes; PE samples in cluster-2 showed early-onset, more severe symptom and worse outcomes.

### Methods for identifying differentially expressed genes (DEGs)

Two widely used tools in R platform, DESeq2^56^ and edgeR,^57^ were applied to determine differential expression genes in different comparisons. The genes with at least 1 count in every samples were used in DESeq2 method and the genes with CPM (Counts Per Million) value larger than 1 in at least one sample were used in edgeR method.

Samples similarity in each group was calculated using all information of expressed genes (raw count of gene in all samples is more than 1) for heat-map, and top 500 genes with the biggest variations were used for PCA analysis. Four comparisons between the clinical subtypes of PE patients were carried out using both DESeq2 and edgeR: **(i)** all PE placental samples (n = 33) versus all normal placental samples (n = 32); **(ii)** late-onset mild PE placental samples (n = 9) versus all normal placental samples; **(iii)** early-onset severe PE placental samples (n = 9) versus all normal placental samples; and **(iv)** late-onset severe PE placental samples (n = 15) versus all normal placental samples (Fig. 2). Adjusted *P*-value in DESeq2 and FDR in edgeR were used together to determine whether a gene is differentially expressed. If adjusted *P*-value ≤ 0.05 of one gene under DESeq2 and FDR ≤ 0.05 under edgeR, the gene was considered as differentially expressed in between two groups of samples. The intersections of differential expression genes from these two methods were taken as the final results of each comparison. Because we only identified a very small number of the differential expression genes in LOMPE, we merge the results of DESeq2 and edgeR as the final result in the comparison of LOMPE samples with normal samples. The numbers of differentially expressed genes identified by the two methods and gene biotypes were showed in **Figure S2A** and **Table S3**.

**Fig. 2.**
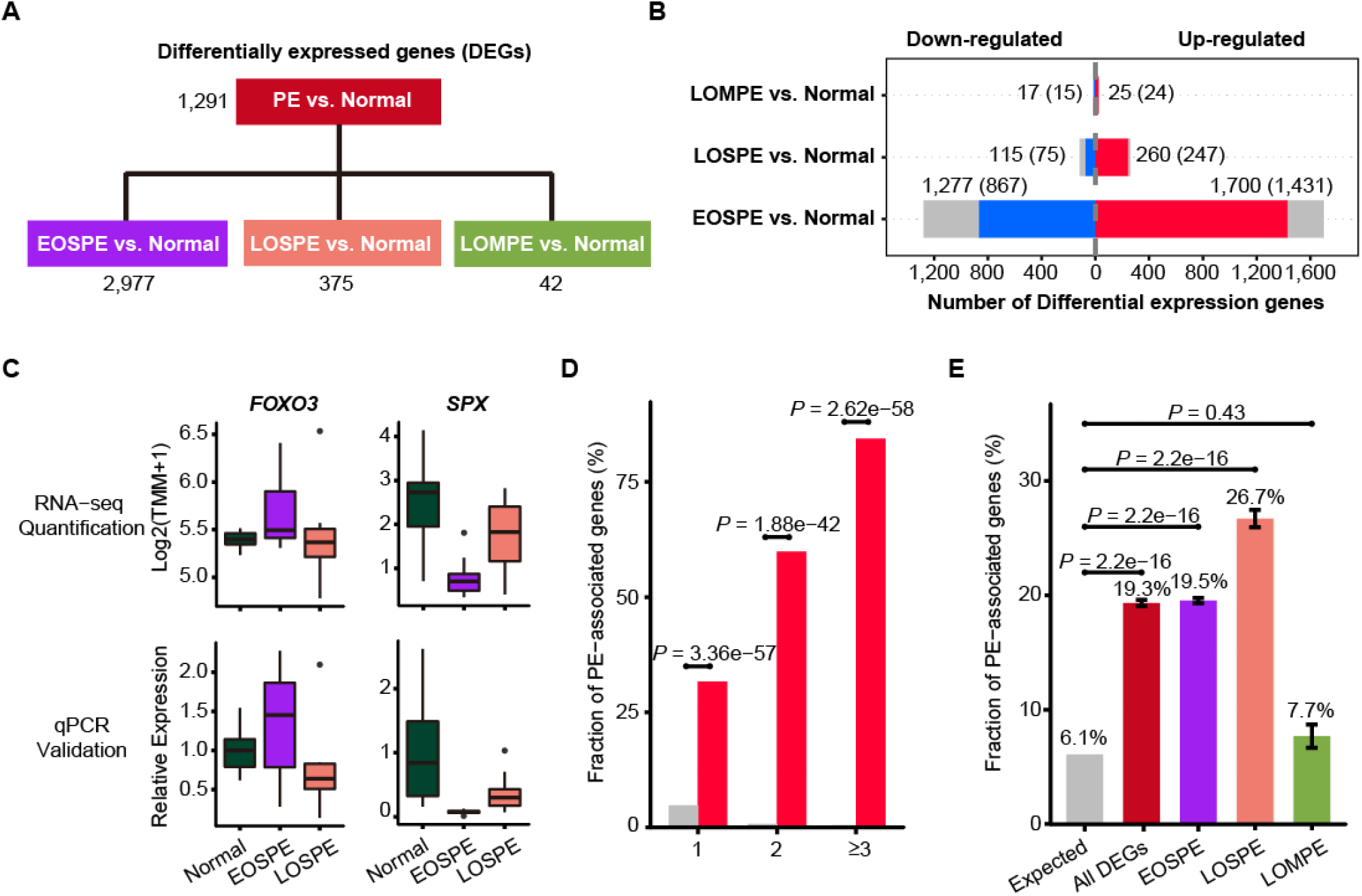
Differentially expressed genes in preeclamptic placentae. **(a)** The differentially expressed genes (DEGs) for each group were identified by comparing data from all PE samples combined or EOSPE, LOSPE and LOMPE samples with normal samples using two methods, DESeq2 and edgeR. **(b)** The red bars represent up-regulated genes, the blue bars represent down-regulated genes and the grey bars non-coding genes. Numbers represent DEGs in each group, and numbers in brackets represent protein-coding DEGs. **(c)** Examples of DEGs verified by qPCR. FOXO3: RNA-seq quantification boxplot at top-left panel and qPCR validation result at bottom-left panel. SPX: RNA-seq quantification boxplot at top-right panel and qPCR validation result at bottom-right panel. **(d)** The enrichment of PE associated genes in DEGs. PE-associated genes were curated from literature (Table EV4). Of the genes that appear once, twice and three or more times in different literature, 31.7%, 59.9% and 84.4% were recovered by our DEGs. The X-axis represents the number of times of a gene reported in the literature, the Y-axis represents the fraction of reported PE-associated genes under each category, with red bars representing ratio of genes that appear once, twice, and three or more times in different literature, and gray bars the expected ratios. **(e)** The enrichment of known PE-associated genes in protein-coding DEGs in EOSPE (2,298 genes), LOSPE (322 genes), LOMPE (39 genes) and all DEGs (2,405 genes). We collected 1,177 PE-associated genes from the literature (Table EV4), and calculated the their enrichment in DEGs of different groups: red bar represents the fraction of PE-associated genes in all DEGs (19.3%, *P* = 2.2e-16), purple bar EOSPE (19.5%, *P* = 2.2e-16), pink bar LOSPE (26.7%, *P* = 2.2e-16%) and light-green bar LOMPE (7.7%, *P* = 0.4261), compared to gray bar which represents the expected ratio (6.1%) of 1,177 PE-associated genes to 19,351 human coding genes (GRCH38.p12 ENSEMBL Gene V93). One-sided Fisher’s exact test was used to compare the nodes kept with the nodes removed. Error bars represent the standard error of the fraction, estimated using bootstrapping method with 100 resamplings.

### The DEGs overlapped with PE-associated genes from the literature

There were a lot of studies to investigate differentially expressed genes compared normal placenta tissues with PE placenta tissues. Due to small sample sizes and undistinguished subtypes of PE samples in previous transcriptomic studies, the differentially expressed genes in PE have not been well defined. To increase the sample sizes, several meta-analyses on reported microarray datasets were published since 2012.^58–63^ In 2015, Kaartokallio et al reported the first RNA-seq study on PE.^46^ To obtain a consensus PE-associated genes, we collected 1,177 genes from literature, including 6 meta-analysis papers,^58–63^ 1 literature-curation paper^64^ and 1 RNA-seq research paper.^46^ These 1,177 genes are PE-associated genes and information for genes were listed in **Table S4**.

### Pathway enrichment analysis for DEGs of EOSPE-only, LOSPE-only and shared genes

The numbers of differentially expressed protein-coding genes are 2,061 in EOSPE-only, 85 in LOPSE-only and 237 in common (Venn diagram in **Figure S3A**). KEGG pathway enrichment analysis was done by clusterProfiler package^54^. The cutoff of *q*-value ≤ 0.2 was chosen to select most significantly enriched pathways (**Figure S3A**). There 39 enriched pathways for EOSPE-only DEGs, 35 for LOSPE-only DEGs, and 7 for shared DEGs.

The pathway enrichment for up-regulated and down-regulated DEGs in both EOSPE and LOPSE were done using the same method (**Figure S3B, 3C**).

### GO terms enrichment analysis for DEGs of EOSPE and LOSPE

Gene Ontology terms, (BP, CC and MF) enrichment analyses were also using clusterProfiler package,^54^ and the *q*-value ≤ 0.05 and adjusted *p*-value ≤ 0.05 were set as cutoff to select most significantly enriched terms. The DEGs of EOSPE were statistically significant enriched in 248 terms of biological process (BP), 51 cellular component (CC) and 16 molecular function (MF). The DEGs of LOSPE were statistically significant enriched 169 terms of BP, 22 of CC and 5 of MF. The overlap of three categories of enriched GO terms are 63 (BP), 4 (CC) and 0 (MF) between DEGs of EOSPE and LOSPE, and most of enriched terms are different in EOSPE and LOSPE (**Figure S3D, 3E and Table S5**).

We used semantic similarity-based method (REVIGO)^65^ to summarize the enriched GO terms in the tree components. For EOSPE, 199 of 248 BP terms were summarized into 14 representative entries. The first representative entry is the nuclear-transcribed mRNA catabolism, containing 55 terms. The second is SRP-dependent co-translational protein targeting to membrane, containing 42 terms. Other entries include stress-activated protein kinase signaling cascade, regulation of macro-autophagy, response to metal ion, estrogen and insulin, and superoxide metabolism. For LOSPE, however, 130 of 169 BP terms were summarized into 11 representative entries. The first entry is positive regulation of cell adhesion, containing 57 terms, and the second entry was response to interferon-gamma, containing 27 terms. Other entries include response to oxygen levels, temperature homeostasis, epithelial cell proliferation, and synapse organization.

The Enriched CC and MF terms in EOSPE and LOSPE also showed considerable difference (**Table S5**). This suggested that besides “canonical pathways” such as HIF-1 signaling pathway, the enriched terms are associated with metabolisms, substrates transport in EOSPE, whereas enriched terms are associated with immune functions in LOSPE.

We found that there are 5 terms of MF enriched in EOSPE DEGs are related to glucose/carbohydrate transport, and 20 terms of BP enriched in EOSPE DEGs are also associated with materials transmembrane transporters, while 17 terms of BP enriched in LOSPE DEGs are associated with transporters.

### Clustering analysis for GO-term-to-gene-bipartite networks for differentially expressed transporter genes of EOSPE and LOSPE

We found that transporter genes were significantly enriched in the DEGs of the EOSPE (*P* = 0.027) compared with all annotated human genes (19,738) in GO, but not significantly enriched DEGs of the LOSPE (*P* = 0.723). There were 205 and 23 transporter genes involved in 397 and 93 GO biological process (GO BP) terms in EOSPE and LOSPE respectively. To summarize the enriched BP terms, we firstly constructed GO-term-to-gene-bipartite networks for EOSPE and LOSPE in Cytoscape using yfiles organic layout. Then we calculated Jaccard score using number of shared genes in the terms, and clustered terms into several clusters based on shared genes. Combining the layout of networks and the results of clustering analysis, 5 and 2 groups of GO BP terms were manually determined for EOSPE and LOSPE, respectively. All enriched BP terms in differentially expressed transporter genes of the EOSPE and the LOSPE were listed in **Table S7**.

### Transcription factors and target genes

To find out the transcription factors that may mostly determine the distinct gene expression patterns in EOSPE and LOSPE, we matched the target genes collected from in 6 databases^66–72^ to the 1,639 reliable human transcription factors (TFs) downloaded from a recent literature.^73^ WE obtained 996 TFs and 16,538 target genes (**Table S8**) and performed TF-target network analysis (Fig. 5 **and Figure S5**).

### Schizophrenia associated genes enriched in the DEGs of the EOSPE and LOSPE

Abnormal intra-uterine environment is an important status of preeclampsia, and our data showed that 90% and 58% babies born by EOSPE and LOSPE patients had FGR/LBW. Recently, a meta-analysis reported that hypertensive disorders of pregnancy may be associated with an increase in the risk of ASD (autism spectrum disorder) and ADHD (attention-deficit/hyperactivity disorder)^74^. Weinberger’s study showed that the set of genes within genomic risk loci for schizophrenia that have interaction with intra-uterine and perinatal complications is highly expressed in placenta^21^. Other studies also demonstrated that preeclampsia was associated with increased risk of offspring schizophrenia.^75^

To investigate the relationship between preeclampsia and schizophrenia, we collected 3,437 schizophrenia-associated genes from classical schizophrenia databases and literature.^76–82^ Since increasing evidences demonstrate the association of preeclampsia with schizophrenia, we want to know if the differentially expressed genes are enriched for schizophrenia associated genes. In our dataset, the protein coding DEGs (2,405, including DEGs from EOSPE, LOSPE and LOMPE), DEGs of EOSPE (2,298) or LOSPE (322) were significant enriched for schizophrenia-associated genes: 20.7% (497/2,405, *P* = 5.0e-5), 20.54% (472/2,298, *P* = 1.4e-4), 22.98% (74/322, *P* = 0.01), compared with the expected 17.8% (the ratio 3,437 schizophrenia associated genes to 19,351 annotated human coding genes). In protein coding DEGs of LOMPE, there is no significant enrichment of schizophrenia associated genes (15.4%, *P* > 0.5) (**Figure S2D**).

## RESULTS

### Subtypes based on clinical characteristics and sample clustering based on RNA-seq data

We collected 32 placentae from 30 normal individuals and 33 placentae from 30 PE patients, which were carefully classified into subtypes based on clinical characteristics (Fig. 1A **and Table S1**). The disease group and normal group showed significant differences in systolic and diastolic pressures, delivery time and baby weights. The early-onset severe PE (EOSPE) group showed significant differences in most of these characteristics compared to the late-onset severe PE (LOSPE) cases. Both EOSPE and LOSPE showed significant differences in these characteristics compared to late-onset mild PE (LOMPE), and the LOMPE group had only moderately elevated blood pressures compared to controls (see **Table S1** for statistics and *P* values, Student’s t test). Notably, the LOSPE patients had significantly elevated C-reactive protein (CRP) levels (*P* = 0.0203 between LOSPE and normal control, *P* = 0.0268 between LOSPE and EOPSE, Student’s *t* test), whereas the other two groups were within the normal range (**Figure S1A**).

Four placental specimens were dissected from the midsection between the chorionic plate and maternal basal plate of each placenta soon after delivery and rapidly preserved in RNAlater® to prevent RNA degradation. RNA-seq was carried out with Illumina Highseq2500. We also performed RNA-seq on two cord-blood samples for computational removal of contamination of blood cells in the placental tissues (**Table S2**). Principal component analysis (PCA) and sample clustering analysis were performed on our dataset, to explore sample features in gene expression levels. The EOSPE samples were clustered together with four LOSPE samples in PCA (Fig. 1B) and in the heatmap (Fig. 1C). The rest LOSPE and all LOMPE samples were mixed with normal samples (Fig. 1B, 1C). After the normal samples were removed from the clustering analysis on the PE samples, we got similar results (**Figure S1B and Table S1**). Looking into the clinical records, we found that the four LOSPE patients whose samples were clustered together with EOSPE samples, showed characteristics similar to those of the EOSPE patients except for gestational age at PE-onset, indicating these four “LOSPE” patients could in fact be EOSPE patients who were diagnosed later than the actual onset time. We extracted the top 50 genes that contributed to the first principal component (PC1), and found 36 known PE-associated genes such as *BHLHE40*, *ENG*, *FLT1*, *HK2*, *HTRA4*, *INHBA*, *LEP*, *NDRG1*, *PAPPA2*, *SASH1*, *SIGLEC6*, *SLC6A8*. The signatures of gene expression that separate EOSPE from the two other clinical subtypes suggest that they may have different underlying molecular mechanisms.

### Distinct gene expression patterns in PE subtypes

To identify groups of genes that may be involved in the heterogeneous pathology of the clinical subtypes, we compared the RNA-seq data of all PE samples or each clinical subtype with the normal controls, using two different methods DESeq2 and edgeR (Fig. 2B), and found a total number of 3,100 differentially expressed genes (DEGs), including 31 microRNAs and 399 lncRNAs (**Table S3**).

We identified 2,977 DEGs (2,298 protein-coding genes) by comparing EOSPE samples with controls, while we only got 1,291 DEGs by comparing all PE samples with controls, suggesting a homogeneity within the EOSPE cases. For the other two subtypes, we found 375 DEGs (322 protein-coding genes) between LOSPE samples and controls, and 42 DEGs (39 protein-coding genes) between LOMPE samples and controls (Fig. 2A **and Table S3**). The large differences between the numbers of DEGs in the three clinical subtypes (EOSPE, LOSPE and LOMPE) indicate that there are different molecular mechanisms underlying the clinical subtypes (Fig. 2B). In general, we found more up-regulated genes than down-regulated genes. The numbers of up-regulated genes are 1.3∼2.3 times higher than those of down-regulated genes in the three comparisons. We carried qPCR to validate some of the differentially expressed genes (Fig. 2C**, Figure S2B, 2C**).

To assess the functional relevance of our dataset, we compared our DEGs with 1,177 PE-associated genes collected from the literature (**Table S4**). Most of the reliable PE-associated genes which appear twice or more in literature are recovered in our dataset (Fig. 2D). Of the 1,177 known PE-associated genes, the DEGs in our dataset (protein-coding genes) recover 31.7% (296/934), 59.9% (88/147) and 84.4% (81/96) of those found once, twice, three or more times in literature. These PE-associated genes show significant enrichment in DEGs (protein-coding genes) of EOSPE and LOSPE: 19.3% (*P* = 2.2e-16, One-sided Fisher’s exact test) in all DEGs, 19.5% (*P* = 2.2e-16, One-sided Fisher’s exact test) in DEGs of EOSPE, 26.7% (*P* = 2.2e-16, One-sided Fisher’s exact test) in DEGs of LOSPE, and 7.7% (*P* = 0.4261, One-sided Fisher’s exact test) in DEGs of LOMPE (Fig. 2E). Because PE has the most significant consequence on developmental brain impairments in offspring, which lead to intellectual disability,^15^ epilepsy,^16^ autism^17–19^ and schizophrenia,^20, 21^ we performed the enrichment analyses of schizophrenia-associated genes in DEGs, and found that the disease-associated genes were significantly enriched (**Figure S2D**). These results suggest that the differentially expressed genes are not only associated with PE, but also with consequence of PE on the offspring in their later life.

### Molecular Pathways involved in PE

The large differences between DEGs observed in EOSPE and LOSPE suggest that these two major subtypes are due to different molecular mechanisms. To find out the associated pathways and molecular functions of the DEGs of these two major subtypes, we performed enrichment analyses on the Kyoto Encyclopedia of Genes and Genomes (KEGG) and Gene Ontology (GO) terms for the protein-coding DEGs of EOSPE (2,298) and LOSPE (322) (Fig. 3). We found that 39 enriched pathways for EOSPE and 35 enriched pathways for LOSPE (**Table S5**). Importantly, the enriched pathways for EOSPE and LOSPE were largely distinct, with only 7 pathways in common. The common pathways included the HIF-1 signaling pathway, focal adhesion, carbon metabolism, glycolysis/gluconeogenesis, biosynthesis of amino acids, adipocytokine and the AGE-RAGE signaling pathway (Fig. 3A **and Table S5**). Most of the enriched pathways of EOSPE were involved with metabolism, such as lysosomal function, AMPK signaling, tight junction pathways, insulin signaling, fatty acid elongation, galactose metabolism, alanine, aspartate and glutamate metabolism pathways (Fig. 3A), many of which are reported to be associated with PE in previous studies (**Table S5**). By contrast, most of the enriched pathways of LOSPE were involved in immune or autoimmune pathways, such as allograft rejection, inflammatory bowel disease, rheumatoid arthritis, HTLV-I infection and tuberculosis pathways (Fig. 3A). The immune genes *HLA-DRB1*, *HLA-DQA1*, *HLA-DRA*, *HLA-DPA1*, *HLA-DPB1* and *HLA-DMB* were up-regulated in LOSPE, but not in EOSPE. These results suggest that the two main subtypes EOSPE and LOSPE have different underlying mechanisms. The subtype EOSPE may be mainly caused by disturbance in metabolism, while the subtype LOSPE by dysfunction of the immune system in the placenta. The metabolic disturbances in EOSPE and the abnormal immune functions in LOSPE may lead to changes in other pathways in the placenta. We obtained similar results in the pathway enrichment analysis for DEGs of EOSPE-only, LOSPE-only and shared DEGs between the two subtypes (**Figure S3A**). We also performed pathway enrichment analysis for up-regulated and down-regulated DEGs respectively, and found that the up-regulated genes and down-regulated genes were involve in different pathways in EOSPE, but no enriched pathway for down-regulated genes in LOSPE (**Figure S3A, 3B and Table S5**).

**Fig. 3.**
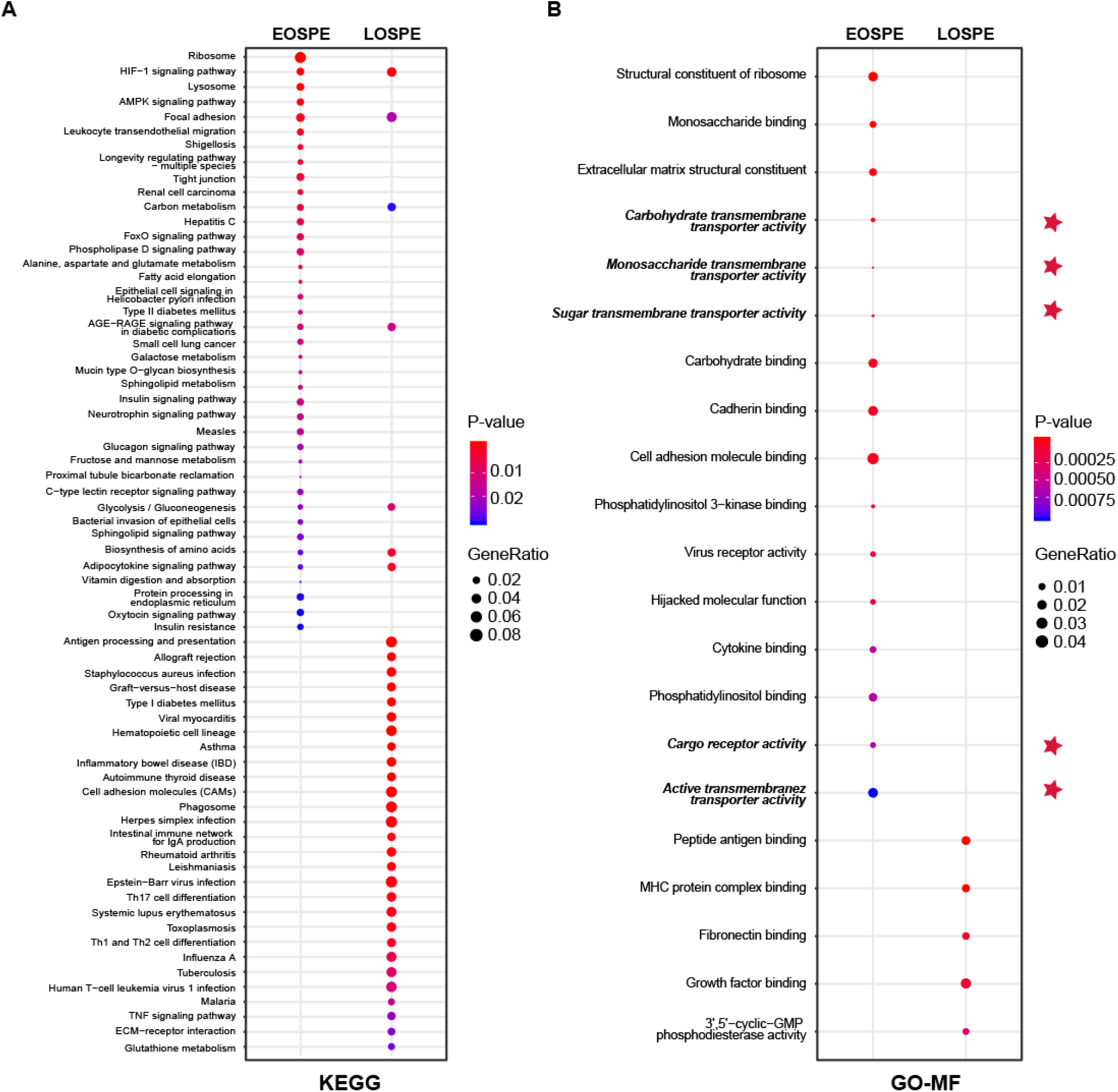
Enrichment of KEGG pathways and GO molecular function (MF) terms with differentially expressed protein-coding genes of EOSPE and LOSPE. **(a)** Enriched KEGG pathways with protein-coding DEGs of EOSPE and LOSPE. **(b)** Enriched GO MF terms with protein-coding DEGs of EOSPE and LOSPE. Dots represent the enriched KEGG pathways or GO MF terms with description of each pathway and term; colors represent scale of *P*-values, the sizes of dots represent ratio of DEGs in corresponding pathways and GO terms, and red stars indicate transporter terms in GO MF. Detailed information is listed in Table S5.

We also performed GO-term enrichment analysis, and found that the DEGs in EOSPE were involved in distinct molecular activities (GO-MF) from those in LOSPE (Fig 3B). We detected significant enrichment of DEGs of EOSPE in 16 molecular function terms of GO, including key molecular activities in constituents of ribosomal structure, differential sugar binding and transmembrane transporter activities, phosphatidylinositol binding, adhesion-molecule binding, virus receptor activities and molecular functions hijacked by virus (Fig. 3B). We detected significant enrichment of DEGs of LOSPE in only 5 molecular function terms of GO, three of which are involved in the immune response, and the other two terms are growth factor binding and 3’,5’-cyclic-GMP phosphodiesterase activity (Fig. 3B). These results are consistent with the results of pathway enrichment analysis we showed above (Fig. 3A): the principal molecular disturbance in EOSPE in the placenta involved metabolism, while immune-function abnormalities predominated in LOSPE. The results of enrichment analyses on biological process (BP) terms and cellular component (CC) terms of GO support the same conclusion (**Figure S3C, 3D and Table S5**).

### Transporter genes involved in PE

Because the DEGs of EOSPE were enriched in many transmembrane transporter activities of GO MF terms (Fig. 3B), we collected 1,554 transporter genes by searching The Gene Ontology (GO) database with key words “transporter”, “carrier” or “substrate exchanging” in order to further look into the dysregulated molecular transportation in the placenta of EOSPE patients (**Table S6**). We found 13.2% (205 of 1,554) transporter genes in DEGs of EOSPE, and only 1.5% (23/1,554) transporter genes in DEGs of LOSPE (Fig. 4B **and Table S6**). Notably, these transporter genes were significantly enriched in the DEGs of the EOSPE (*P* = 0.027, One-sided Fisher’s exact test), but not in LOPSE (*P* = 0.723, One-sided Fisher’s exact test) (Fig. 4B).

**Fig. 4.**
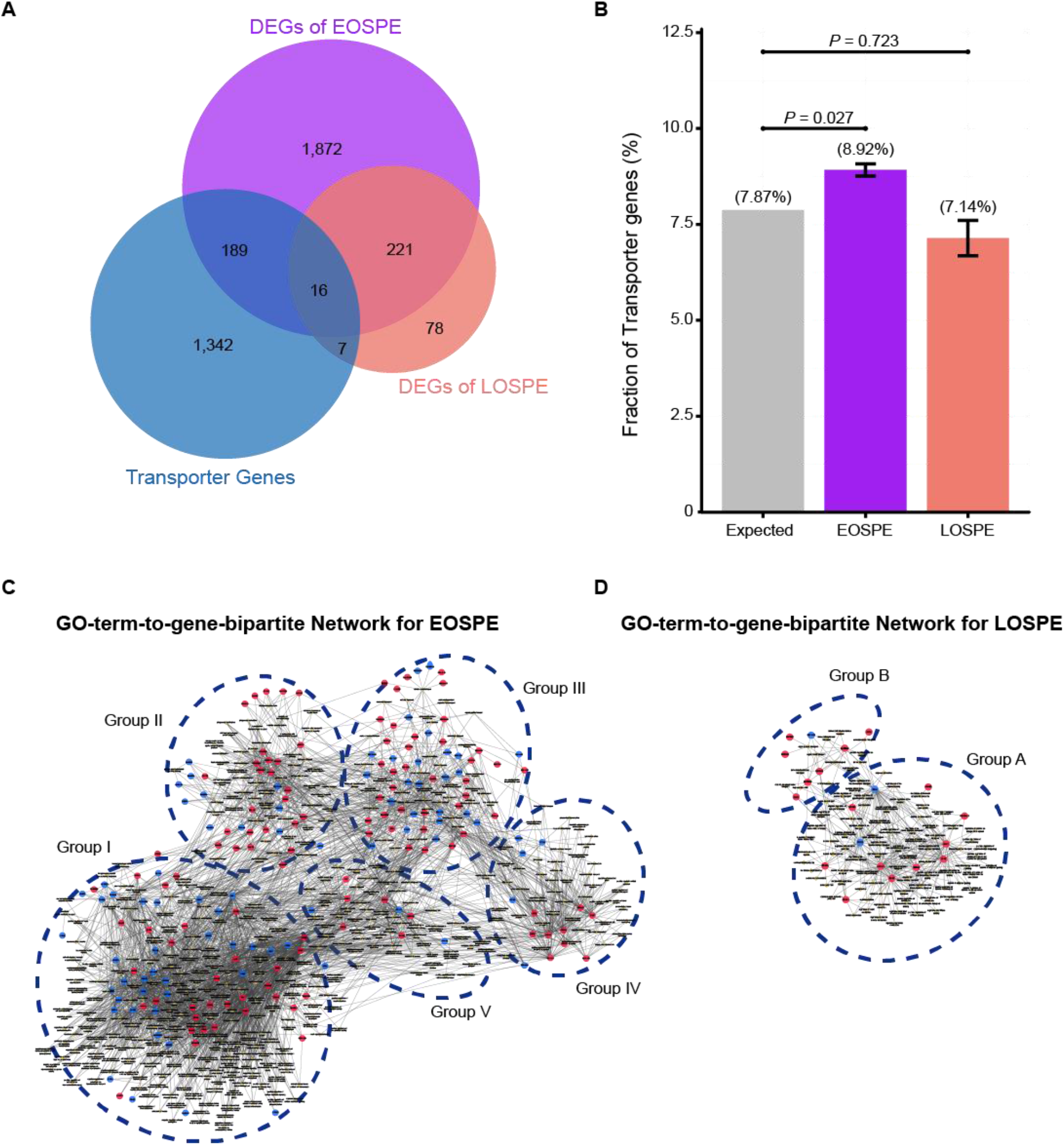
Functional classification of differentially expressed transporter genes. **(a)** Venn diagram for the overlapping genes between the transporter genes and DEGs. Of the identified transporter genes, 205 were differentially expressed in EOSPE and 23 genes in LOSPE. **(b)** Fraction of transport genes in DEGs. The transporter genes were collected from Gene Ontology. Gray bar represents the expected fraction (7.87%) of transporter genes in all GO annotated genes (19,738), the fraction of transporter genes in the DEGs of EOSPE samples (8.92%) (purple bar), the DEGs of LOSPE (7.14%) (pink bar). The transporter genes were significantly enriched in the DEGs of EOSPE, but not in LOSPE. One-sided Fisher’s exact test was used to compare the nodes kept with the nodes removed. Error bars represent the standard error of the fraction, estimated using bootstrapping method with 100 resamplings. **(c)** Gene-GO term bipartite network for enriched GO BP terms and DEGs in EOSPE. Blue dashed circles show 5 biological process clusters involved in different transportation processes of substrates and materials. Group I: ion channels; group II: macro-autophagy; group III: transmembrane transportation of nonlipid nutrients; group IV: transmembrane transportation of lipids; group V in transportation of hormone. **(d)** Gene-GO term bipartite network for enriched GO BP terms and DEGs in LOSPE. The group A: ion channels; group B: transportation of chloride, carboxylic acid, cAMP and anion. Blue dashed circles show 2 biological process clusters. Red nodes represent up-regulated transporter genes and blue nodes down-regulated transporter genes, and yellow diamond nodes represent GO BP terms. The edges indicate the GO BP terms the genes belong to.

The 205 transporter genes in DEGs of EOSPE were enriched in 397 GO biological process (BP) terms (Adjusted *P*-value ≤ 0.05, hypergeometric test from ClusterProfiler Package). For these genes, we constructed a GO-term-to-gene bipartite network for all enriched BP terms, which contained 5 groups of functionally related genes (Fig. 4C **and Table S7**). Group I contains genes involved in different ion channels, some of which were previously reported to be associated with PE, e.g. *ATP1A1*, *CACNA1D*, *CALM1*, *EHD3*, *FGF12*, *KCNE1*, *KCNH5*, *NOS1AP*, *PDE4D*, *SCN1B*, *SCN4B* and *SRI* (**Table S7**); group II contains genes involved in macro-autophagy, which was also previously proposed to be involved in PE,^83, 84^ such as some up-regulated genes *ATP6V1B1*, *ATP6V1C2*, *ATPV1D1*, *ATP6V0A1*, *ATP6V0E1*, and *ATP6V0D1;* group III, genes in transmembrane transportation of hexose, fatty acids, amino acids, vitamins, organic anions and mitochondrial components; group IV, genes in transmembrane transportation, storage or homeostasis of lipids, sterols, cholesterol and phospholipids, and group V, genes in transportation of hormone, response of cAMP and drug, secretion of insulin and protein. (Fig. 4C). The detailed information of all the five groups is included in the Supplemental material (**Table S7**). We also performed pathway enrichment analysis for these 205 differentially expressed transporter genes in EOSPE and found that the genes were enriched in 34 pathways (**Figure S4 and Table S7**), which are consistent with the five functional groups in GO-term analysis (Fig. 4C). Some of these pathways are known to be involved in PE, such as oxidative phosphorylation,^40^ ABC transporter pathway^85^ and circadian entrainment pathway.^86^

The 23 transporter genes in the DEGs of LOSPE were enriched in 93 GO biological process terms (Adjusted *P*-value ≤ 0.05, hypergeometric test from ClusterProfiler Package). We also constructed a GO-term-to-gene-bipartite network for these 93 terms, which contained 2 groups of functionally related genes (Fig. 4D). The group A contains 15 genes involved in ion channels, and group B only contains 8 genes involved in transportation of chloride, carboxylic acid, cAMP and anion.

The placenta serves as a molecular barrier between maternal and fetal circulation, and transmembrane transporters play a key role in the placental exchange function. On one hand, the fetus receives nutrients from the mother, such as carbohydrates, lipids, amino acids, vitamins and minerals for adequate development and growth, and on the other hand, the fetus can drain waste products via efflux processes into the maternal circulation.^87^ The dysregulation of the quantity, density and activity of membrane transporters could impact the nutrient supply and consequently affect fetal development. We consistently observed low birth weight (LBW) and fetal growth restriction (FGR) in 90% (9 of 10) of the babies born by EOSPE patients and 53% (8 of 15) of the babies born by LOSPE patients (**Table S1**). The weights of babies in EOSPE group were significantly lower than those of babies in LOSPE group (**Table S1**).

### Transcription factors involved in PE

We want to know if the distinct differential gene expression patterns in EOSPE and LOSPE might be due to the dysregulation of a set of transcription factors. We used 1,639 reliable human transcription factors (TFs)^73^ from literature to search 6 databases^66–72^ for their targets, and built a regulatory network containing 996 TFs and 16,538 target genes (Fig. 5A). These 996 TFs cover 65.6% (105/160) of the TFs in the DEGs of EOSPE, such as *LTF*, *BCL6*, *ARNTL*, *NFKB1*, *PPARG* and *ATF3* (**Table S8**), which target to 81.9% (1,882/2,298) of DEGs in EOSPE. They also cover 81.5% (22/27) of TFs in DEGs of LOSPE, such as *FOSL1*, *FOXO1*, *STAT3*, *YBX1*, *DBP* and *SOX15* (**Table S8**), which target 67.1% (216/322) of DEGs in LOSPE (Fig. 5B, 5C **and Table S8**).

**Fig. 5.**
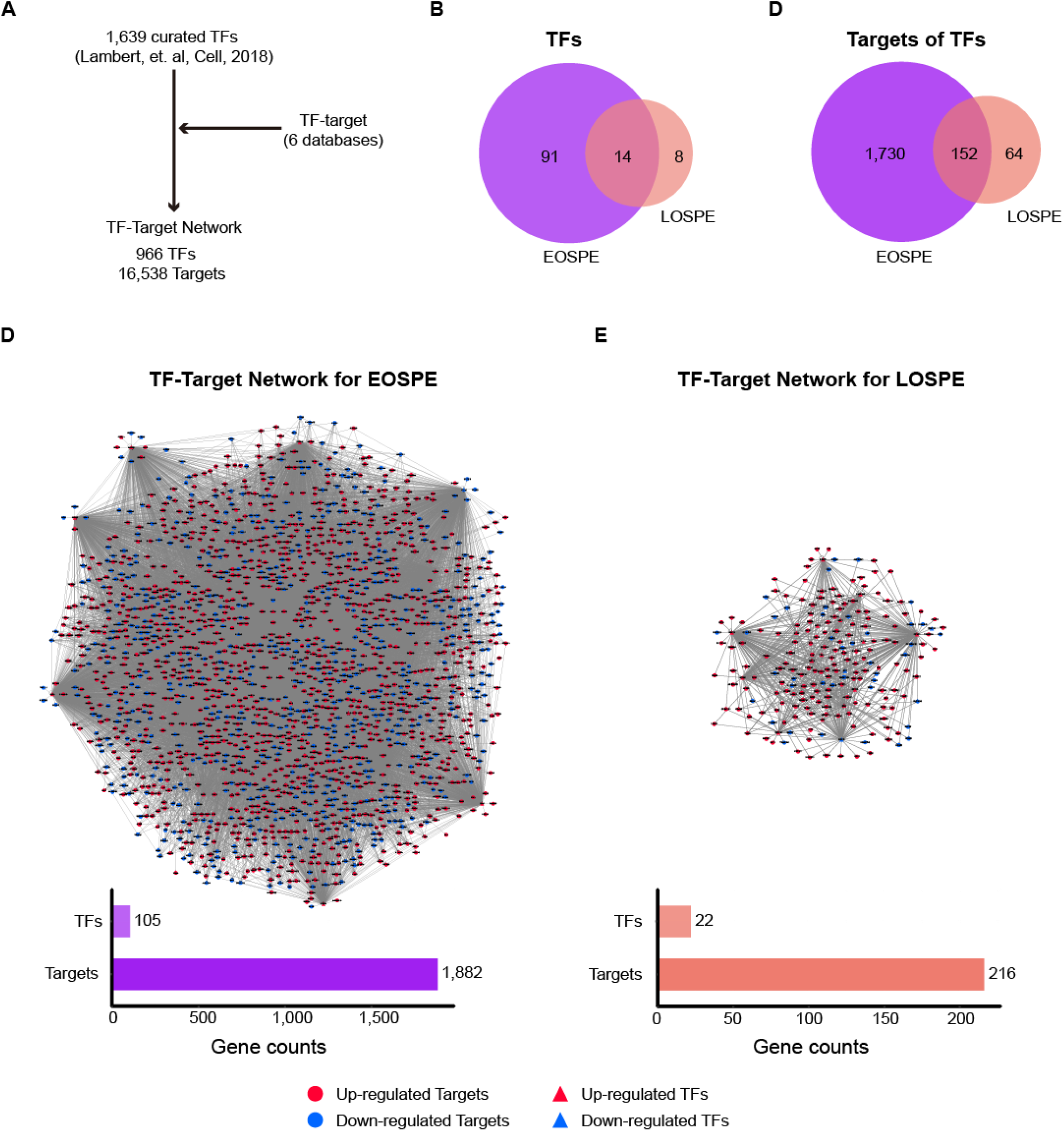
TF-target networks for EOSPE and LOSPE. **(a)** Workflow for extracting TF-target network for EOSPE and LOSPE. We collected TF-target data from 6 databases,^66–72^ and then matched the 1,639 reliable human transcription factors (TFs)^73^ to their targets and constructed a TF-target network containing 996 TFs and 16,538 target genes. **(b)** The comparison between curated TFs and DEGs in EOSPE and LOSPE: 91 TFs differentially expressed in EOSPE only; 8 TFs in LOSPE only; 14 TFs shared by both. **(c)** The comparison between TF targets of the DEGs in EOSPE and LOSPE: 1,882 (81.9%) DEGs in EOSPE regulated by the 105 TFs, and 216 (67.1%) DEGs in LOSPE by the 22 TFs. **(d)** TF-target network for EOSPE. The bar chart shows the numbers of differentially expressed TFs (105) and their differentially expressed targets (1,882) in EOSPE. **(e)** TF-target network for LOSPE. The bar chart shows the numbers of differentially expressed TFs (22) and their differentially expressed targets (216) in LOSPE. Red circles represent up-regulated genes, blue circles down-regulated genes, red triangles represent up-regulated TFs and blue triangles down-regulated TFs. Edges with arrows represent regulation of TFs to their target genes.

These TF-to-target regulation relationships were exemplified as regulatory networks (Fig. 5D, 5E), suggesting that altered expression in a small number of TFs may lead to a large number of DEGs in PE. We performed pathway enrichment analysis on the two regulatory networks and found that 18 pathways were involved in EOSPE and 47 pathways in LOSPE (**Table S8**), which are consistent with the enriched pathways for the differentially expressed genes in EOSPE and LOSPE (Fig. 4). Some key PE-related pathways were affected by dysregulated TFs as shown in the two example modules involved in AMPK and FoXO signaling pathways (**Figure S5A, 5B**) in EOSPE and two other example modules in PI3K-Akt signaling pathway and immune function pathways in LOSPE (**Figure S5C, 5D**).

## DISCUSSION

Patients with PE display a wide range of symptoms. Early-onset PE patients usually have severe symptoms; among the late-onset PE patients, however, some show severe symptoms, while others are only mildly affected. Although the clinical characteristics of PE can be measured, the underlying molecular bases for the clinical subtypes are not known. Anecdotal studies on placentae of PE patients point to some abnormal molecular processes, such as defective angiogenesis,^88, 89^ hypoxia^90–92^ and inflammation,^39, 93, 94^ but these abnormalities do not unambiguously define the different molecular pathways involved in the clinical subtypes. Additional approaches are needed to identify the genes underlying the differential pathogenesis of the PE subtypes. Numerous microarrays on transcriptomes of placentae^37–44^ and a recent RNA-seq study^46^ did not identify distinct molecular processes associated with the clinical subtypes, probably due to small sample sizes and inadequate comparisons between the grouped samples. The other transcriptomic study on the PE focuses on the maternal side, the uterine decidua.^95^ In order to systematically search for genes involved in clinical PE subtypes, we dissected samples from the mid-section placental tissues between the chorionic and maternal basal surfaces of placentae from patients with classical presentation of preeclampsia, which were grouped into three clinical subtypes: early-onset severe PE (EOSPE), late-onset severe PE (LOSPE) and late-onset mild PE (LOMPE) (**Table S1**). Our identification of the distinct molecular processes involved in the clinical subtypes was based on initial classification and comparison of RNA-seq data of the well classified samples. Because we noticed that the blood could not be completely washed away from the placental tissues, which might be one factor affecting previous transcriptomic analyses, we performed RNA-seq on whole cord-blood cells and removed cord-blood contamination by computational method (**Table S2**).

In the principal component analysis (PCA) and sample clustering analysis, the EOSPE samples grouped together, separate from the clusters of LOSPE and LOMPE samples (Fig. 1B). We carried out comparisons between the placental transcriptomes of the three clinical subtypes and normal samples (EOSPE vs. normal, LOSPE vs. normal and LOMPE vs. normal) to identify the differentially expressed genes in the three clinical subtypes. Surprisingly, EOSPE had 2,977 differentially expressed genes (2,298 protein coding genes), while LOSPE had only 375 (322 protein coding genes), indicating different molecular mechanisms for these two subtypes (Fig. 2A); both data sets, however, showed significant enrichment of previously reported PE-associated genes (Fig. 2C). Moreover, LOMPE had only 42 differentially expressed genes that showed no enrichment for the previously reported PE-associated genes (Fig. 2C), suggesting a lesser degree of involvement of genetic factors in the pathogenesis of mild PE. Most interestingly, we found distinct molecular processes involved in the two major subtypes of PE: metabolisms and related activities in the EOSPE and immune activities in LOSPE (Fig. 3). These PE-associated pathways were regulated by small number of transcription factors (Fig. 5), which might be the driver genes during the abnormal development of the placenta of PE patients.

As the placenta is the supporting system for a developing embryo, the widespread gene expression change in placentae of PE patients might have profound effects on the development of the fetuses and the health of the offspring in later life. Accumulating evidences point to association of preeclampsia with brain developmental disorders, including intellectual disability,^15^ epilepsy,^16^ autism^17–19^ and schizophrenia.^20, 21^ It might be partially due to the placenta-derived factors that affects the development of the brain and thus leads to complex brain disorders.^21, 74^ Ursini et al found that, for some loci of schizophrenia, the association between genetic risk and schizophrenia is affected by complicated pregnancies, and the gene set within these loci are differentially expressed in placentae from complicated in comparison with normal pregnancies^21^. In order to investigate to what extent the differentially expressed genes in PE may be also involved in schizophrenia, we collected 3,437 schizophrenia-associated genes from classical schizophrenia databases and literature, and found that the schizophrenia-associated genes are significantly enriched in DEGs of EOSPE (*P* = 1.4e-4, One-sided Fisher’s exact test) and LOSPE (*P* = 0.01, One-sided Fisher’s exact test) (**Figure S2D**).

In conclusion, this study provides molecular-level evidences that EOSPE and LOSPE are two different subtypes with distinct underlying molecular mechanisms while LOMPE may not be due to placental factors. Expression of metabolism-related genes is significantly affected in EOSPE, whereas in LOSPE it preferentially involves the expression of immune-related genes. Most notably, the transporter genes dysregulated in EOSPE may cause abnormal molecular exchange between maternal and fetal circulation, leading to fetal growth restriction. The TF-target networks demonstrate that only a small number of dysregulated transcription factors may serve as driving genes, causing differential expression of a larger number of genes in EOSPE and LOSPE. Such distinct underlying mechanisms suggest that potentially more effective therapies could be developed to target the different subtypes at their molecular levels.

## Supporting information

Supplementary Information (Supplementary Figures included)

## ACKNOWLEDGEMENTS

We thank the patients for their generous support for this study. This work was supported by Science and Technology Programme of Guangdong Province (No. 2017B020227010 and No. 2015B050501006) and National Natural Science Foundation of China (No.31771434, No. 81671466, No 81471464 and No. 81771609).

