## Supplementary Information (Supplementary Figures included) for "Distinct Molecular Processes in Placentae Involved in Two Major Subtypes of Preeclampsia"

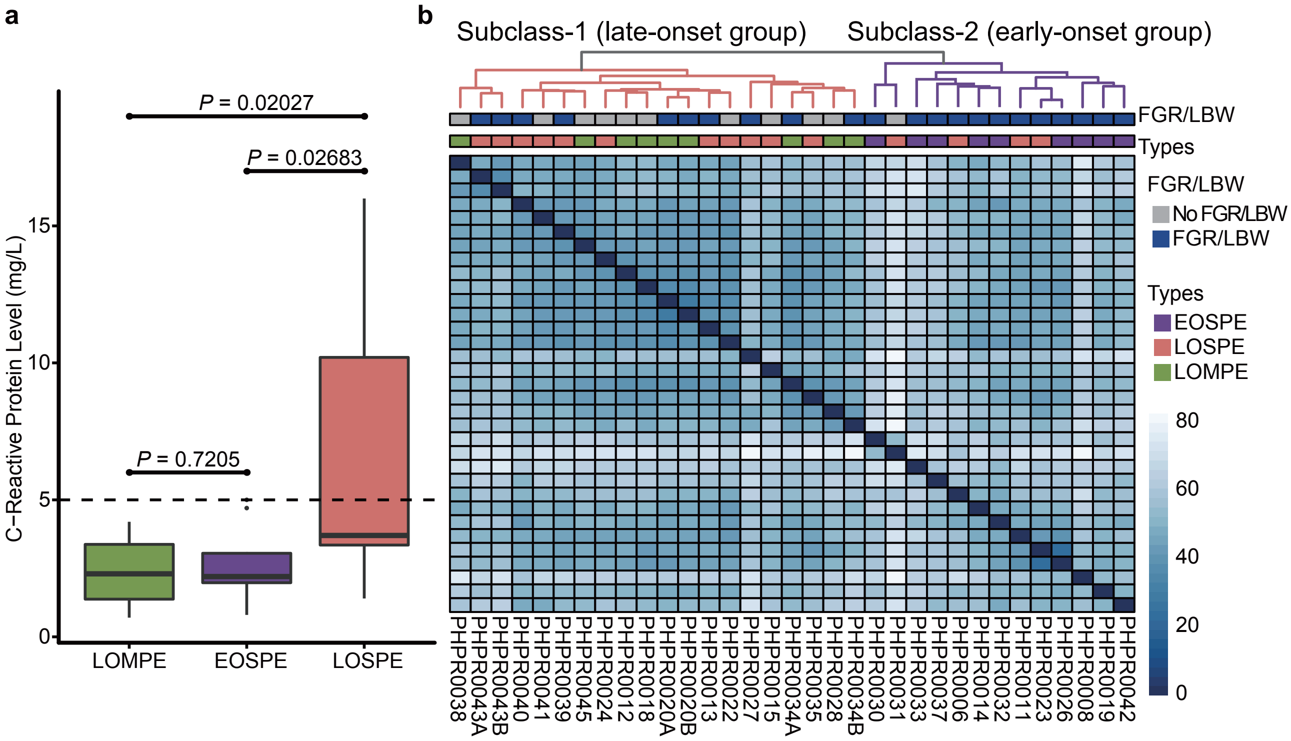


**Figure S1. C-reactive protein levels in clinical subtypes and sample clustering analysis based on RNA-seq data of early-onset severe, late-onset severe and mild samples. (a)** Box-plot of c-reactive protein levels in LOMPE, EOSPE and LOSPE. The normal range of c-reactive protein levels is 0-5mg/L. **(b)** Heatmap of sample-sample distance clustering for disease samples (late-onset severe, mild and early-onset severe samples) using Ward.D. Two clear subclasses were observed, where most of late-onset severe together with mild samples clustered as subclass-1 and early-onset severe together with four late-onset severe samples clustered as subclass-2.

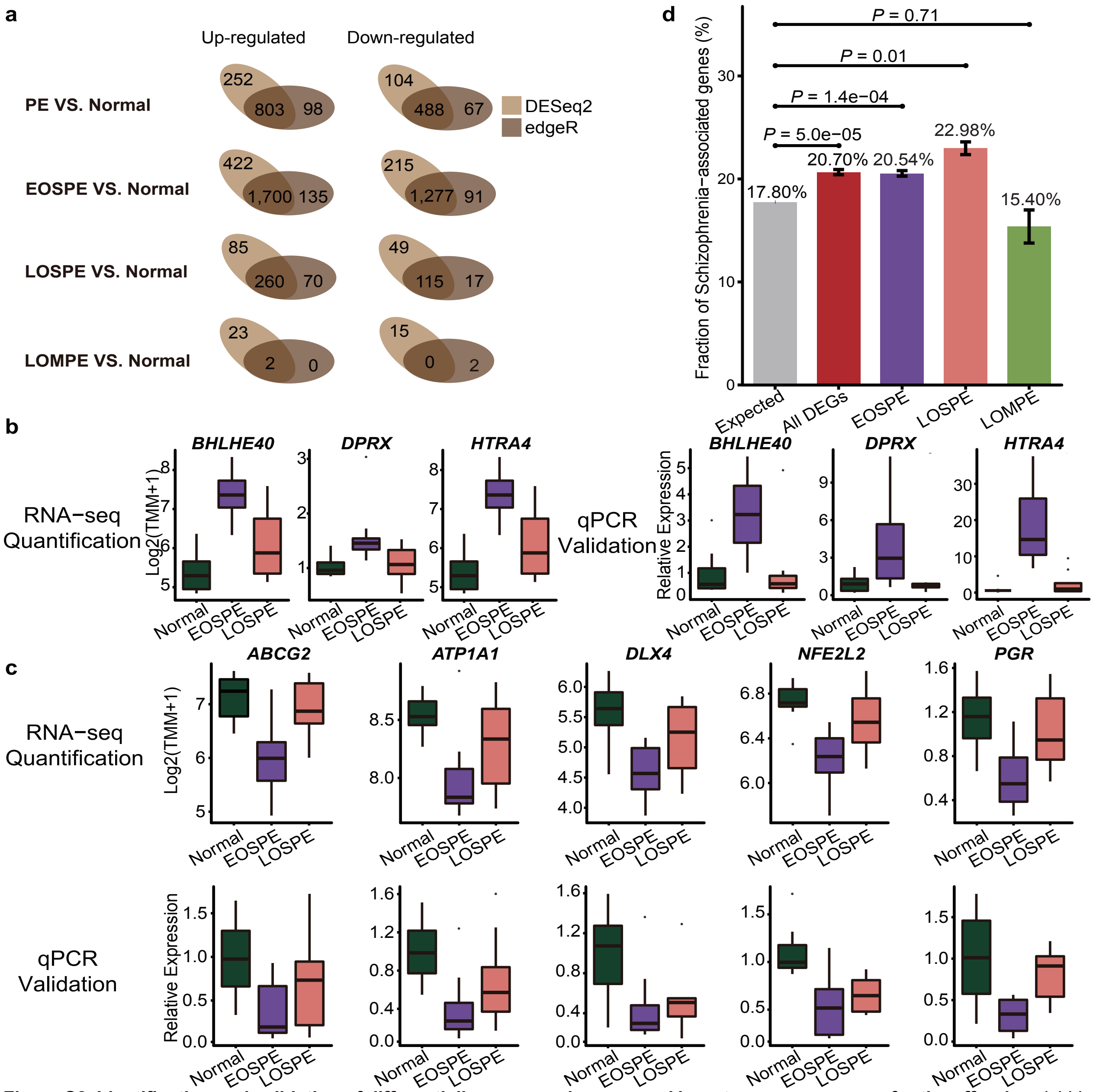


**Figure S2. Identification and validation of differentially expressed genes, and long-term consequence for the offspring. (a)** Venn diagrams for identifying differentially expressed genes using DESeq2 and edgeR methods in four comparison groups. Left Venn diagrams are number of up-regulated genes detected by DESeq2 and edgeR and right Venn diagrams are number of down-regulated genes detected by DESeq2 and edgeR. The overlapped genes were taken as the final differentially expressed genes in each comparison group. Light brown represents method of DESeq2; dark brown method of edgeR. **(b)** Box-plots for four up-regulated genes identified from RNA-seq and q-PCR validation. **(c)** Box-plots for six down-regulated genes identified from RNA-seq and q-PCR validation. **(d)** Schizophrenia-associated genes were enriched in DEGs of preeclampsia. The schizophrenia associated genes collected from classical schizophrenia databases and literature showed significantly enrichment in DEGs in all PE samples combined, EOPSE or LOPSE, whereas no enrichment in DEGs in LOMPE. Gray bar represents the expected fraction (17.8%) of schizophrenia associated genes in all protein-coding genes (19,351), the fraction of schizophrenia-associated genes in DEGs in all PE samples (20.7%), the DEGs of EOSPE (20.54%), LOSPE (22.98%) and LOMPE (15.4%) were presented as red, purple, pink and light green bars. The *P*-value was calculated using the Fisher's exact test, and error bars represent the standard error of the fraction, estimated using bootstrapping with 100 resamplings.

a

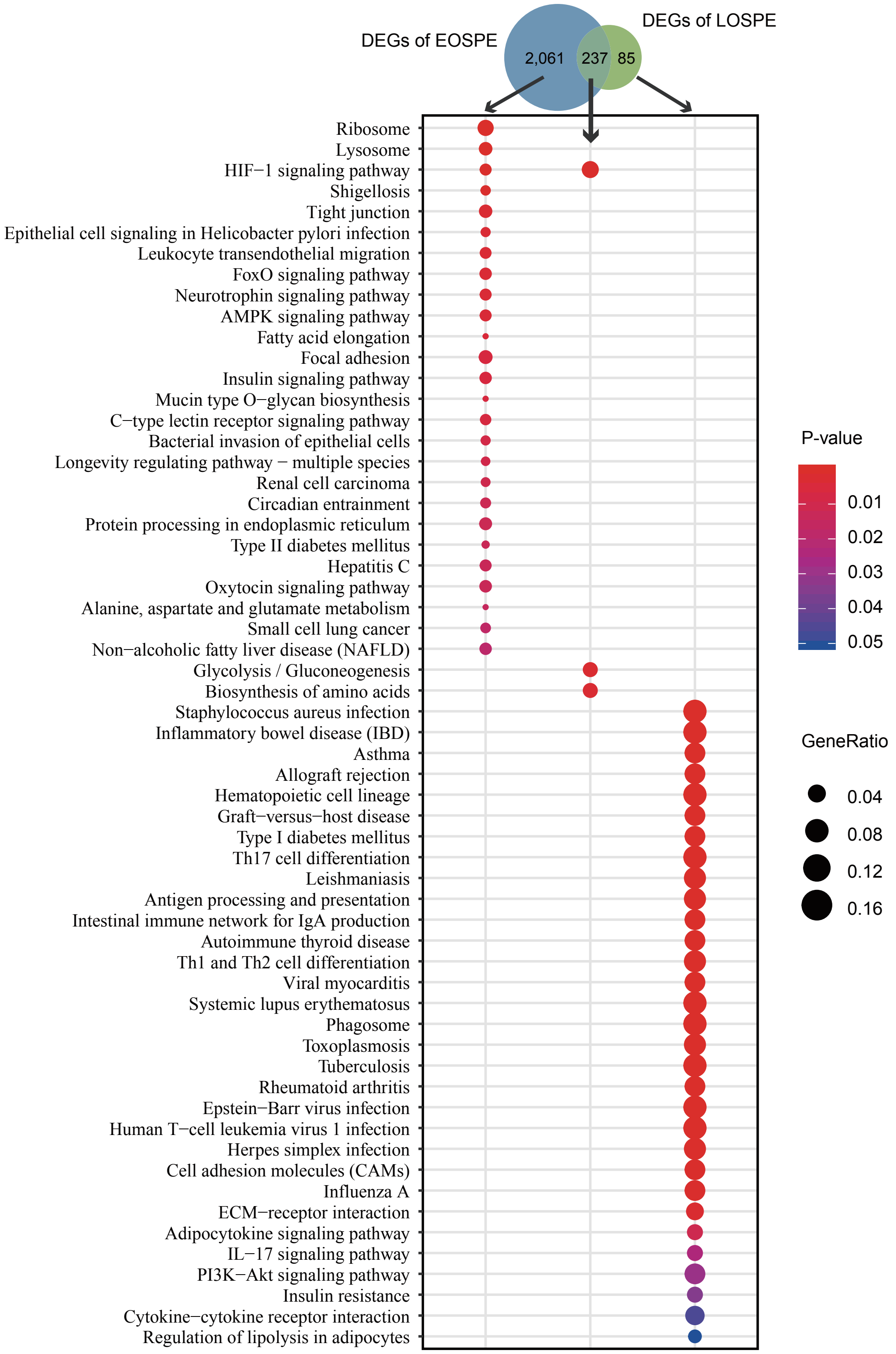


**Figure S3. Functional enrichment analysis of differentially expressed genes. (a)** Enrichment of DEGs in KEGG pathways for EOSPE-only, common or LOSPE-only DEGs. Venn-diagram shows the overlaps between DEGs of EOSPE and LOSPE. **(b)-(c)** Enrichment of DEGs in KEGG pathways for up- and down-regulated DEGs in the EOSPE **(b)** and LOPSE **(c)**. **(d)** Enrichment of DEGs in GO BP terms for DEGs of EOSPE and LOSPE. **(e)** Enrichment of DEGs in GO CC terms for DEGs of EOSPE and LOSPE. Dot colors indicate enrichment *P*-values and dot sizes represent gene ratios in the enriched pathways.

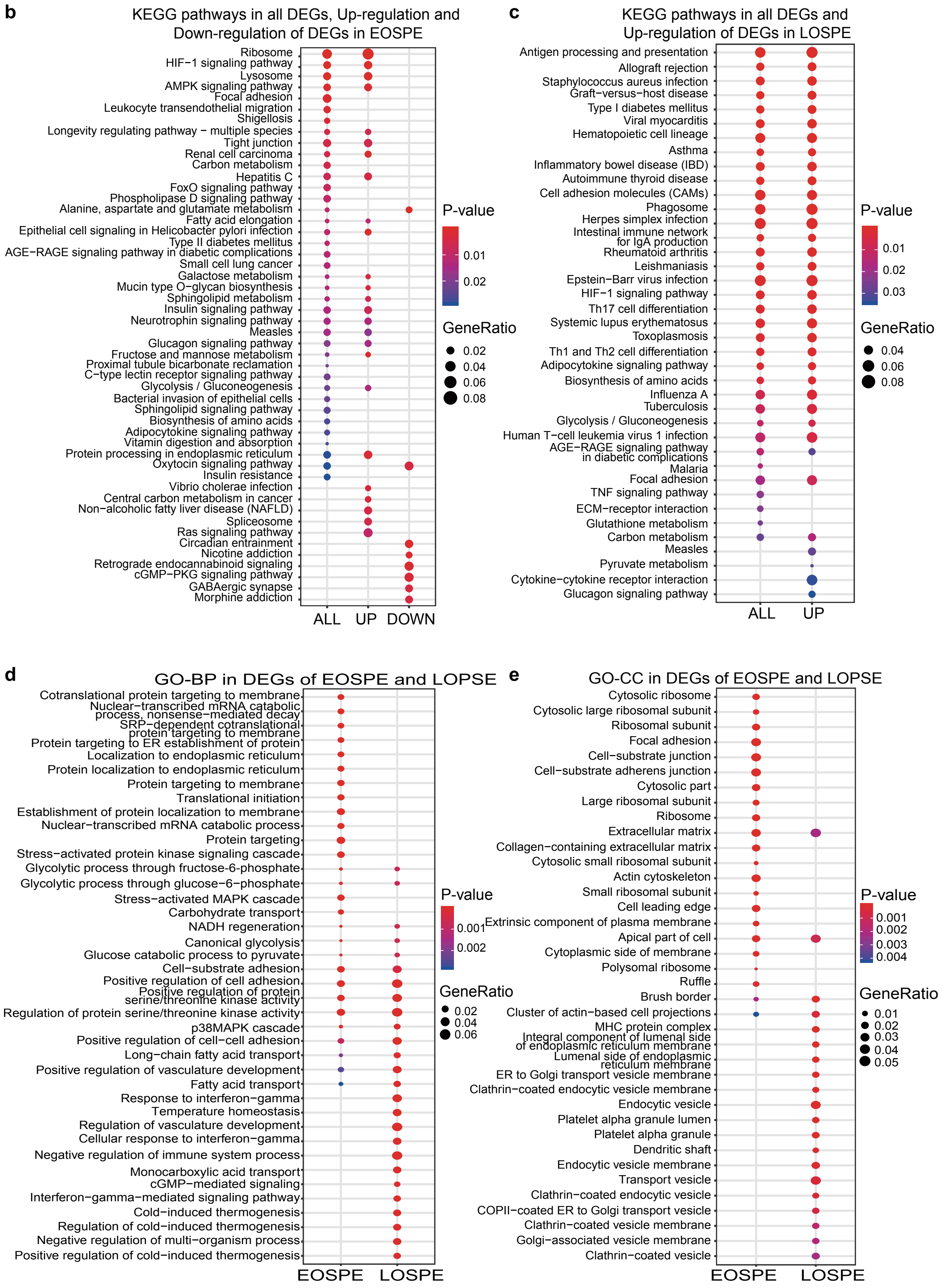
b
KEGG pathways in all DEGs, Up-regulation and Down-regulation of DEGs in EOSPE
Ribosome
HIF−1 signaling pathway
Lysosome
AMPK signaling pathway
Focal adhesion
Leukocyte transendothelial migration
Shigellosis
Longevity regulating pathway − multiple species
Tight junction
Renal cell carcinoma
Carbon metabolism
Hepatitis C
FoxO signaling pathway
Phospholipase D signaling pathway
Alanine, aspartate and glutamate metabolism
Fatty acid elongation
Epithelial cell signaling in Helicobacter pylori infection
Type II diabetes mellitus
AGE−RAGE signaling pathway in diabetic complications
Small cell lung cancer
Galactose metabolism
Mucin type O−glycan biosynthesis
Sphingolipid metabolism
Insulin signaling pathway
Neurotrophin signaling pathway
Measles
Glucagon signaling pathway
Fructose and mannose metabolism
Proximal tubule bicarbonate reclamation
C−type lectin receptor signaling pathway
Glycolysis / Gluconeogenesis
Bacterial invasion of epithelial cells
Sphingolipid signaling pathway
Biosynthesis of amino acids
Adipocytokine signaling pathway
Vitamin digestion and absorption
Protein processing in endoplasmic reticulum
Oxytocin signaling pathway
Insulin resistance
Vibrio cholerae infection
Central carbon metabolism in cancer
Non−alcoholic fatty liver disease (NAFLD)
Spliceosome
Ras signaling pathway
Circadian entrainment
Nicotine addiction
Retrograde endocannabinoid signaling
cGMP−PKG signaling pathway
GABAergic synapse
Morphine addiction
ALL
UP
DOWN
P-value
0.01
0.02
GeneRatio
0.02
0.04
0.06
0.08
c
KEGG pathways in all DEGs and Up-regulation of DEGs in LOSPE
Antigen processing and presentation
Allograft rejection
Staphylococcus aureus infection
Graft−versus−host disease
Type I diabetes mellitus
Viral myocarditis
Hematopoietic cell lineage
Asthma
Inflammatory bowel disease (IBD)
Autoimmune thyroid disease
Cell adhesion molecules (CAMs)
Phagosome
Herpes simplex infection
Intestinal immune network for IgA production
Rheumatoid arthritis
Leishmaniasis
Epstein−Barr virus infection
HIF−1 signaling pathway
Th17 cell differentiation
Systemic lupus erythematosus
Toxoplasmosis
Th1 and Th2 cell differentiation
Adipocytokine signaling pathway
Biosynthesis of amino acids
Influenza A
Tuberculosis
Glycolysis / Gluconeogenesis
Human T−cell leukemia virus 1 infection
AGE−RAGE signaling pathway in diabetic complications
Malaria
Focal adhesion
TNF signaling pathway
ECM−receptor interaction
Glutathione metabolism
Carbon metabolism
Measles
Pyruvate metabolism
Cytokine−cytokine receptor interaction
Glucagon signaling pathway
ALL
UP
P-value
0.01
0.02
0.03
GeneRatio
0.04
0.06
0.08
d
GO-BP in DEGs of EOSPE and LOPSE
Cotranslational protein targeting to membrane
Nuclear−transcribed mRNA catabolic process, nonsense−mediated decay
SRP−dependent cotranslational protein targeting to membrane
Protein targeting to ER establishment of protein
Localization to endoplasmic reticulum
Protein localization to endoplasmic reticulum
Protein targeting to membrane
Translational initiation
Establishment of protein localization to membrane
Nuclear−transcribed mRNA catabolic process
Protein targeting
Stress−activated protein kinase signaling cascade
Glycolytic process through fructose-6-phosphate
Glycolytic process through glucose−6−phosphate
Stress−activated MAPK cascade
Carbohydrate transport
NADH regeneration
Canonical glycolysis
Glucose catabolic process to pyruvate
Cell−substrate adhesion
Positive regulation of cell adhesion
Positive regulation of protein serine/threonine kinase activity
Regulation of protein serine/threonine kinase activity
p38MAPK cascade
Positive regulation of cell−cell adhesion
Long−chain fatty acid transport
Positive regulation of vasculature development
Fatty acid transport
Response to interferon−gamma
Temperature homeostasis
Regulation of vasculature development
Cellular response to interferon−gamma
Negative regulation of immune system process
Monocarboxylic acid transport
cGMP−mediated signaling
Interferon−gamma−mediated signaling pathway
Cold−induced thermogenesis
Regulation of cold−induced thermogenesis
Negative regulation of multi−organism process
Positive regulation of cold−induced thermogenesis
EOSPE
LOSPE
P-value
0.001
0.002
GeneRatio
0.02
0.04
0.06
e
GO-CC in DEGs of EOSPE and LOPSE
Cytosolic ribosome
Cytosolic large ribosomal subunit
Ribosomal subunit
Focal adhesion
Cell−substrate junction
Cell−substrate adherens junction
Cytosolic part
Large ribosomal subunit
Ribosome
Extracellular matrix
Collagen−containing extracellular matrix
Cytosolic small ribosomal subunit
Actin cytoskeleton
Small ribosomal subunit
Cell leading edge
Extrinsic component of plasma membrane
Apical part of cell
Cytoplasmic side of membrane
Polysomal ribosome
Ruffle
Brush border
Cluster of actin−based cell projections
MHC protein complex
Integral component of lumenal side of endoplasmic reticulum membrane
Lumenal side of endoplasmic reticulum membrane
ER to Golgi transport vesicle membrane
Clathrin−coated endocytic vesicle membrane
Endocytic vesicle
Platelet alpha granule lumen
Platelet alpha granule
Dendritic shaft
Endocytic vesicle membrane
Transport vesicle
Clathrin−coated endocytic vesicle
COPII−coated ER to Golgi transport vesicle
Clathrin−coated vesicle membrane
Golgi−associated vesicle membrane
Clathrin−coated vesicle
EOSPE
LOSPE
P-value
0.001
0.002
0.003
0.004
GeneRatio
0.01
0.02
0.03
0.04
0.05

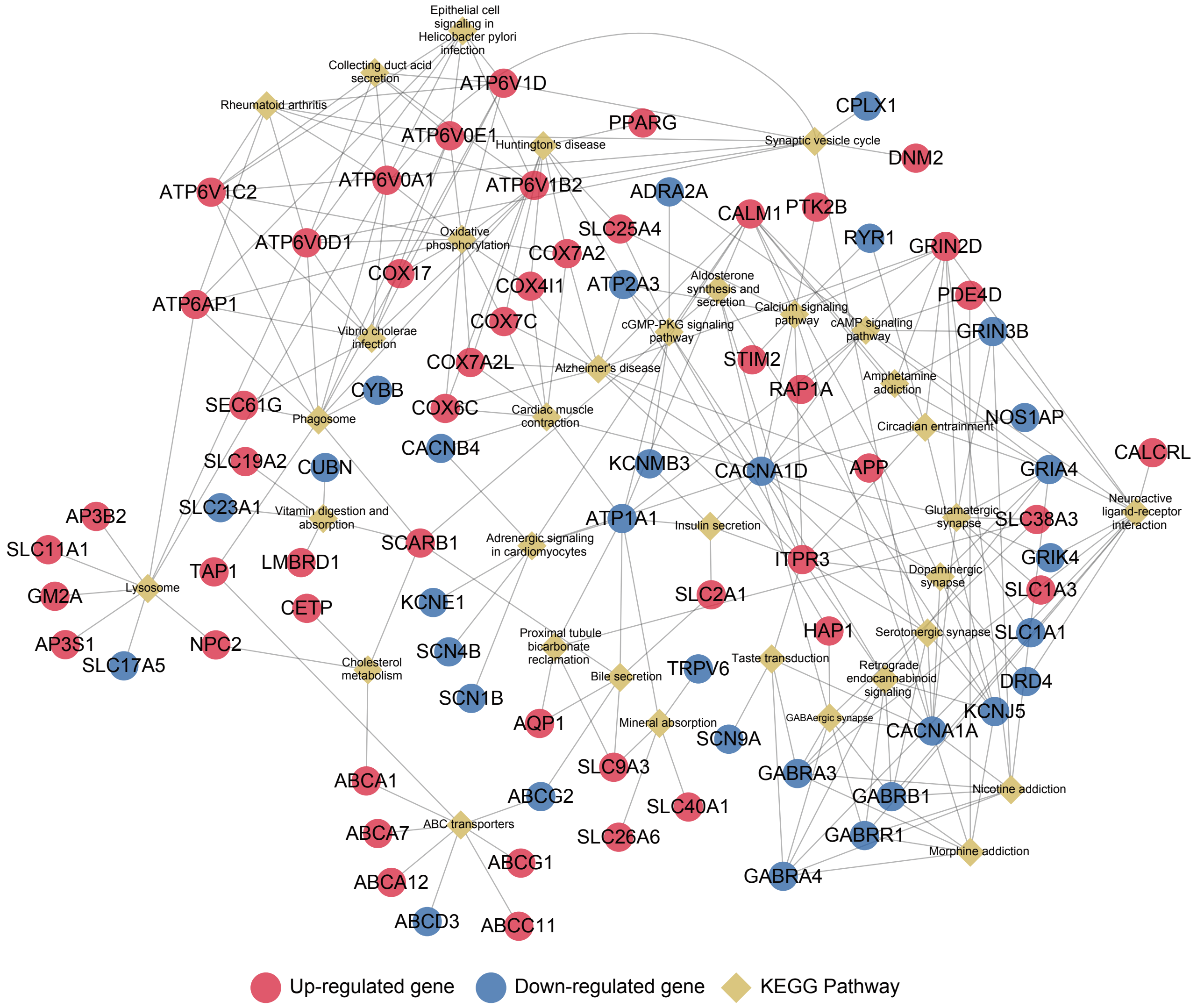


**Figure S4. Gene-pathway bipartite network of DEGs and the enriched KEGG pathways in EOSPE.**
Red nodes represent up-regulated genes and green nodes are down-regulated genes in EOSPE; and yellow diamond nodes represent names of KEGG pathways. The edges connect the DEGs to the KEGG pathways the genes belong to.

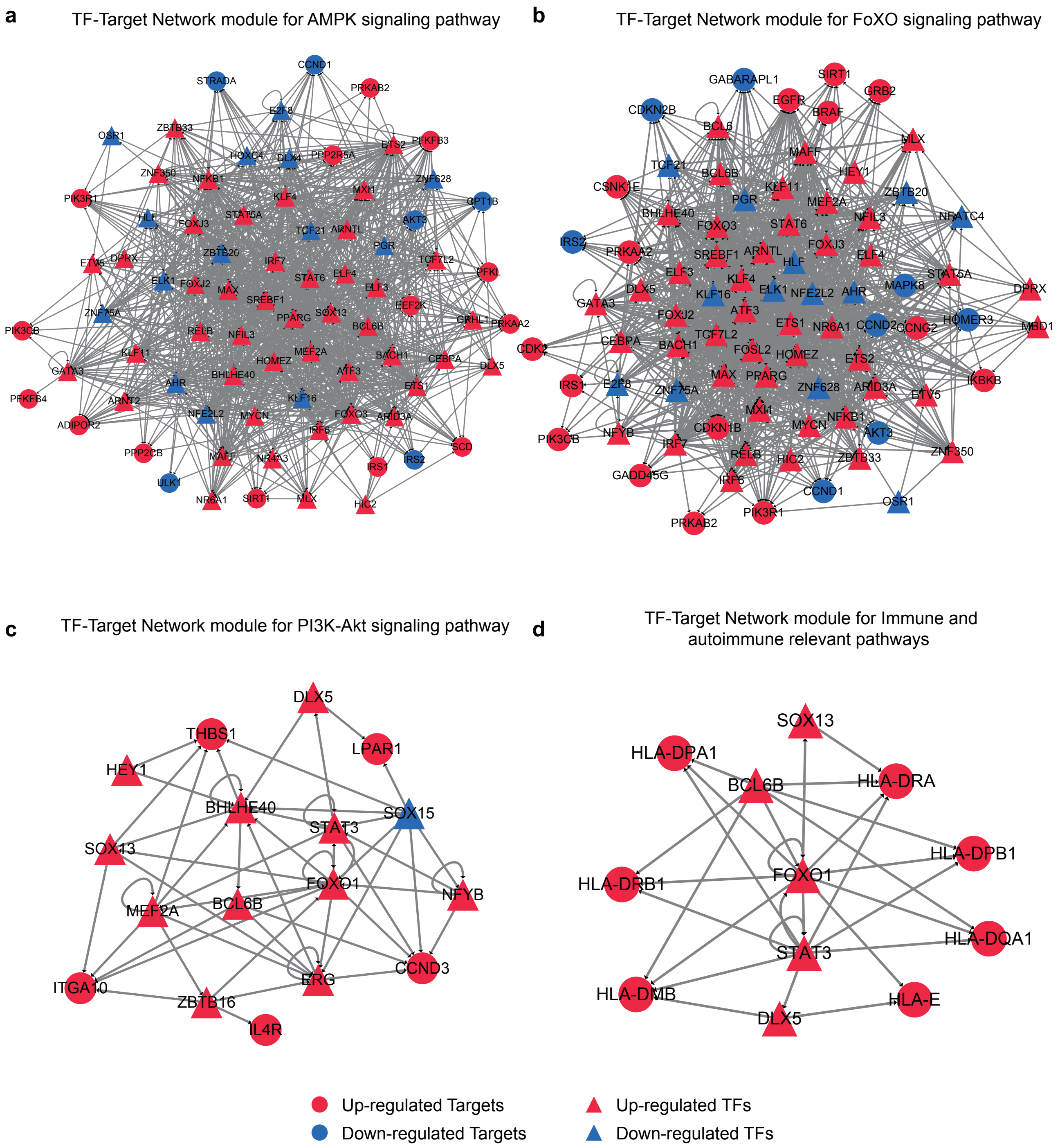


**Figure S5. TF-target network modules for DGEs in EOSPE and LOSPE.** **(a)** TF-target network module for AMPK signaling pathway in EOSPE. **(b)** TF-target network module for FoXO signaling pathway in EOSPE. **(c)** TF-target network module for PI3K-Akt signaling pathway in LOSPE. **(d)** TF-target network module for immune and autoimmune signaling pathway in LOSPE. All genes in TF-target network modules are differentially expressed genes in EOSPE and LOSPE, respectively. The TF targets are members of the pathways.

***Supplementary Tables***

Tables are provided in excel files.

**Table S1. Clinical information.**

Sheet1: Clinical information comparisons among groups of patients with EOSPE, LOSPE or LOMPE and normal subjects;

Sheet2: Clinical information comparison between subclass-1 and subclass-2 PE sample groups.

**Table S2. Raw counts of RNA-seq data.**

Sheet1: Raw counts of each gene for two cord blood samples;

Sheet2: Raw counts of each gene for 65 placental samples before and after removing blood contamination.

**Table S3. Differentially expressed genes (DEGs).**

Sheet1: Gene biotypes and direction of difference of DEGs;

Sheet2: DEGs of All PE;

Sheet3: DEGs of EOSPE;

Sheet4: DEGs of LOSPE;

Sheet5: DEGs of LOMPE;

Sheet6: Q-PCR validation.

**Table S4. PE-associated genes curated from literature.**

Sheet1: Literature information, number of PE-associated genes and datasets used in these studies;

Sheet2: PE-associated genes with times found in the literature and the overlap with DEGs of EOSPE, LOPSE and LOMPE.

**Table S5. Enriched Pathways and GO terms.**

Sheet1: Enriched KEGG pathways in the DEGs of the EOSPE and LOSPE;

Sheet2: Enriched KEGG pathways in the DEGs of the EOSPE-only, LOSPE-only and shared genes in Sheet1;

Sheet3: Enriched KEGG pathways in the up- and down-regulated DEGs of the EOSPE and up-regulated DEGs of the LOSPE;

Sheet4: Enriched GO BP terms;

Sheet5: Enriched GO CC terms;

Sheet6: Enriched GO MF terms;

Sheet7: Representative term groups in BP, CC and MF.

**Table S6. Transporter genes.**

Sheet1: Information for 1,554 transporter genes downloaded from GO and overlapped genes in the DEGs of the EOSPE and the LOSPE;

Sheet2: Information for differentially expressed transporter genes in the EOSPE and the LOSPE.

**Table S7. Enriched GO terms and pathways for transporter genes in DEGs of EOSPE and the LOSPE.**

Sheet1: Information in the GO-term-to-gene-bipartite networks of EOPSE and LOPSE;

Sheet2: Enriched 397 BP terms table in EOSPE;

Sheet3: Enriched 93 BP terms table in LOPSE;

Sheet4: Enriched KEGG pathways in differentially expressed transporter genes of the EOSPE.

**Table S8. Transcription factors and regulatory network.**

Sheet1: Information for 1,639 transcription factors (TFs) and the overlap with the DEGs of the EOSPE and the LOSPE. The first six columns of this table were from Additional file 2: Table S1 of Lambert's study [73];

Sheet2: TF-target regulatory networks for EOSPE. The first column of table is TF and the second column is target regulated by TF;

Sheet3: TF-target regulatory networks for LOSPE. The first column of table is TF and the second column is target regulated by TF;

Sheet4: Enriched biological pathways in TF-target regulatory networks of EOSPE and LOSPE obtained using ReactomeFI plugin of Cytoscape;

Sheet5: TFs involved in enriched pathways.

**Table S9. Tools, databases and primer sequences.**

Sheet1: Tools for analyses;

Sheet 2: Databases;

Sheet 3: Primer sequences for qPCR.
